# A retrospective evaluation of automated optimization of deep brain stimulation parameters

**DOI:** 10.1101/393900

**Authors:** Johannes Vorwerk, Andrea A. Brock, Daria N. Anderson, John D. Rolston, Christopher R. Butson

**Affiliations:** Scientific Computing & Imaging (SCI) Institute, University of Utah, Salt Lake City, UT, USA; Institute of Electrical and Biomedical Engineering, UMIT - University for Health Sciences, Medical Informatics and Technology, Hall in Tirol, Austria; Department of Neurosurgery, Clinical Neurosciences Center, University of Utah, Salt Lake City, UT, USA; Department of Biomedical Engineering, University of Utah, Salt Lake City, UT, USA; Department of Psychiatry, University of Utah, Salt Lake City, UT, USA; Department of Neurology, University of Utah, Salt Lake City, UT, USA

**Keywords:** neuromodulation, essential tremor, computational modeling, thalamus, ventral intermediate nucleus

## Abstract

**Objective:** We performed a retrospective analysis of an optimization algorithm for the computation of patient-specific multipolar stimulation configurations employing multiple independent current/voltage sources. We evaluated whether the obtained stimulation configurations align with clinical data and whether the optimized stimulation configurations have the potential to lead to an equal or better stimulation of the target region as manual programming, while reducing the time required for programming sessions.

**Methods:** For three patients (five electrodes) diagnosed with essential tremor, we derived optimized multipolar stimulation configurations using an approach that is suitable for the application in clinical practice. To evaluate the automatically derived stimulation settings, we compared them to the results of the monopolar review.

**Results:** We observe a good agreement between the findings of the monopolar review and the optimized stimulation configurations, with the algorithm assigning the maximal voltage in the optimized multipolar pattern to the contact that was found to lead to the best therapeutic effect in the clinical monopolar review in all cases. Additionally, our simulation results predict that the optimized stimulation settings lead to the activation of an equal or larger volume fraction of the target compared to the manually determined settings in all cases.

**Conclusions:** Our results demonstrate the feasibility of an automatic determination of optimal DBS configurations and motivate a further evaluation of the applied optimization algorithm.

## 1. Introduction

Deep brain stimulation (DBS) has been established as a treatment for movement disorders, such as Parkinson’s disease and essential tremor [24, 66], and is currently being evaluated as a treatment for a variety of other neurological disorders. Depending on a patient’s diagnosis and symptoms, DBS leads are implanted in specific anatomical targets in the brain during stereotactic surgery. For the patients in our study, the ventrointermediate nucleus (VIM) was chosen as target, which is a common choice to treat essential tremor [9, 45, 37].

A few weeks after implantation of the DBS lead, an initial programming session is performed. During this session, a comprehensive monopolar review is performed in an attempt to find optimal monopolar stimulation settings for the specific patient. Parameters that can be varied are active lead contact(s), stimulation voltage, pulse width, and pulse frequency. For each contact of the DBS lead, the voltage thresholds at which therapeutic or side effects occur for cathodic-phase-first charge-balanced pulses (hereafter referred to as cathodic stimulation) are determined. This monopolar review may require several hours of programming time [33, 44]. If the treatment results are not satisfactory, an exploration of alternative stimulation configurations, such as bipolar or anodic stimulation, may be performed [23, 62, 36, 56]. For most patients, multiple follow-up programming sessions are necessary to refine the stimulation parameters, leading to a significant time effort for both patients and their caregivers. The time expended for the manual programming of DBS patients is expected to grow exponentially with the emerging use of DBS leads that have an increased number of contacts than the previously standard quadripolar leads in combination with implanted pulse generators (IPG) that offer multiple independent current sources. Such stimulation setups are promising better treatment outcomes due to their improved field-shaping abilities [11, 61, 17, 48, 60, 67].

To decrease programming times and to fully capitalize on the possibilities of these novel lead models, different research groups have presented algorithms for an automatic determination of optimal DBS parameters, such as stimulation pattern, strength, etc. [18, 19, 20, 68, 47, 4]. Whereas these algorithms have been positively evaluated in simulation studies, few data exist evaluating the computationally predicted stimulation settings against clinical data.

In this study, we evaluate for the first time automatically optimized configurations for multipolar cathodic stimulation against patient data. These stimulation configurations make use of multiple independent current/voltage sources, i.e., multiple contacts are concurrently stimulating as cathodes at different voltages, and the IPG is used as return electrode. We apply the algorithm proposed by Anderson et al. [4], who evaluated it for leads with both cylindrical and segmented contacts in a simulation study, and compare the determined stimulation patterns to the results of the monopolar review for multiple patients with VIM DBS for essential tremor.

We expanded this optimization algorithm to additionally enforce the efficiency of the stimulation, so that activation of contacts that do not stimulate the target region efficiently is suppressed. To constrain the results of the optimization algorithm to settings that do not cause side effects in practice, we ensured that the optimized stimulation settings did not exceed the side-effect thresholds obtained during the monopolar review. The iterative procedure applied in this study to maximize the stimulation strength while avoiding stimulation settings that cause side effects can similarly be applied in clinical practice, and can potentially replace the time-consuming determination of therapeutic and side-effect thresholds for each contact during the monopolar review.

We performed our study for patients implanted with quadrupolar leads. The main goal of this study is a first, general evaluation of the applied optimization algorithm with regard to the future application for newly available leads with eight or more segmented contacts and IPGs providing multiple independent voltage/current sources [57, 58, 1]. The IPGs implanted with quadripolar leads do not yet support multiple independent voltage/current sources, so that the optimization results obtained in this study can currently not be directly applied in practice. Instead, we compare the optimization results to those of the monopolar review. We find that the optimization algorithm assigns the highest voltage to the contact that was found to lead to the best therapeutic effect in all cases. We further evaluate the predicted target activation and predicted power consumption and find that the increase in predicted target activation exceeds the change in predicted power consumption by more than 50 percentage points in four of five cases.

## 2. Material and methods

### 2.1. Patient cohort and imaging

We obtained data from three DBS patients who were diagnosed with essential tremor. All patients gave written informed consent and all procedures were approved by the independent research board (IRB) of the University of Utah (#44402). Patient details are listed in Table 1. All patients were implanted with Medtronic 3387 leads. For the evaluation of the optimization results, we refer to these patients as PXY, where X is the patient ID between 1 and 3, and Y the stimulation site, i.e., either L for left or R for right.

**Table 1:** Patient overview. *t*_lead_ indicates the time between the initial programming session and lead placement, *t*_IPG_ the time between the initial programming session and IPG placement (w = weeks, d = days).

| | Sex | Age | Hemisphere | ET impairment (off medication) | $t_{\text{lead}}$ | $t_{\text{IPG}}$ |
| --- | --- | --- | --- | --- | --- | --- |
| P1 | W | 53 | Bilateral | upper extremities | 2 w | 3 d |
| P2 | M | 64 | Left | upper extremities, disability of right hand | 5.5 w | 3.5 w |
| P3 | M | 71 | Bilateral | upper extremities, progressive | 5.5 w | 5.5 w |

For all patients, preoperative MP2RAGE T1-weighted (T1w-) scans (voxel size = 1 × 1 × 1 mm, FOV = 255 × 255 × 176 mm) were acquired on a 3T MR scanner (MAGNETOM Prisma 3.0 T, Siemens Healthcare, Erlangen, Germany). Additionally, multi-shell diffusion spectrum (DS-) MRI scans with 64 directions for each b-value (voxel size 1.49 × 1.49 × 1.5 mm, FOV = 250 × 250 × 139.5 mm, b-values = 700, 2000, 3000 s/mm^2^) and volumes with flat diffusion gradient (b = 0 s/mm^2^) for both regular and inverted phase encoding direction were acquired. The DSI volumes were corrected for eddy currents, head motions, and susceptibility artifacts using ACID (http://www.diffusiontools.com; Mohammadi et al. 43, Ruthotto et al. 53) and resampled to an isotropic voxel size of 1 × 1 × 1 mm.

Computed tomography (CT) images were acquired (voxel size = 0.75 × 0.75 × 1 mm) on an AIRO Mobile CT system (Brainlab AG, Feldkirchen, Germany) to reconstruct the lead positions.

### 2.2. DBS programming notes

During each initial programming session, we recorded the voltages, pulse widths, and frequencies at which therapeutic benefit, i.e., no tremor was observed, or side effects occurred for each contact (Table 2). We noted the contact used in the patient’s primary program that was determined to provide the best therapeutic effect during the monopolar review. In some cases, the treating neurologist provided the patient with an alternative program that could be selected using the patient’s personal programmer. Such second-line, alternative contacts were also noted. We further evaluated the observed side effects to identify the brain region whose stimulation caused them. The observed side effects were tingling or pain of arm, leg, trunk, or face (assigned to Vc), speech changes or difficulties (assigned to IC), and induced ataxia (unclear, possibly cerebello-rubrospinal fibers; Reich et al. 51).

**Table 2:** Results of monopolar review. Side-effect thresholds, therapeutic thresholds, pulse width (PW), frequency (FRQ). “SE region” indicates the brain region that was associated with the observed side effects, and “?” marks side effects that could not clearly be assigned to one region (see Section 2.3). “o” marks the contact used in the patient’s primary program, which was determined to provide the best therapeutic effect during the monopolar review. “x” marks the contact used in the patient’s alternative program.

| Patient # | | | C0 | C1 | C2 | C3 | PW [μs] | FRQ [Hz] |
| --- | --- | --- | --- | --- | --- | --- | --- | --- |
| P1R | Side effect | [V] | 0.6 | 0.8 | 0.8 | 3.0 | 60 | 185 |
|  | SE region |  | Vc | Vc | Vc | Vc |  |  |
|  | Therapeutic | [V] | - | - | - | 2.5 |  |  |
|  | Clinically used |  |  |  |  | o |  |  |
| P1L | Side effect | [V] | 0.5 | 1.0 | 1.6 | 1.8 | 60 | 185 |
|  | SE region |  | Vc | Vc | Vc | Vc |  |  |
|  | Therapeutic | [V] | - | 0.8 | 1.5 | 1.6 |  |  |
|  | Clinically used |  |  | x | o |  |  |  |
| P2L | Side effect | [V] | 0.8 | 1.8 | 1.8 | 1.6 | 90 | 180 |
|  | SE region |  | Vc | Vc, ? | Vc | Vc |  |  |
|  | Therapeutic | [V] | 0.7 | 1.5 | - | - |  |  |
|  | Clinically used |  |  | o |  |  |  |  |
| P3R | Side effect | [V] | 0.5 | 0.5 | 1.8 | 2.3 | 60 | 180 |
|  | SE region |  | Vc | Vc | Vc | Vc |  |  |
|  | Therapeutic | [V] | - | - | 1.5 | 1.6 |  |  |
|  | Clinically used |  |  |  | o | x |  |  |
| P3L | Side effect | [V] | 0.7 | 2.8 | 3.4 | 4.0 | 60 | 180 |
|  | SE region |  | Vc | IC, Vc | IC, Vc | IC |  |  |
|  | Therapeutic | [V] | - | 2.0 | 3.0 | 3.8 |  |  |
|  | Clinically used |  |  | o | x |  |  |  |

Given that only (mono- and multipolar) cathodic stimulation with the IPG as the return electrode was considered in both the monopolar review and the computation of optimized stimulation settings, we indicate only the contact and the absolute value of the voltage.

### 2.3. Selection of target and avoidance regions for the optimization algorithm

To apply the optimization algorithm, target regions for which the stimulation is maximized and regions of avoidance for which the stimulation is kept below a certain threshold to minimize side effects have to be selected. In the definition of the algorithm, we refer to this threshold as the sensitivity threshold. We chose the VIM as the target region and the ventralis caudalis (Vc) and the internal capsule (IC) as avoidance regions (see, e.g., Figure 1, Krauth et al. 38, Bakay 7, Figure 11.2), because stimulation of Vc or IC can lead to parasthesias or dysarthria and motor contractions, respectively [46].

**Figure 1:**
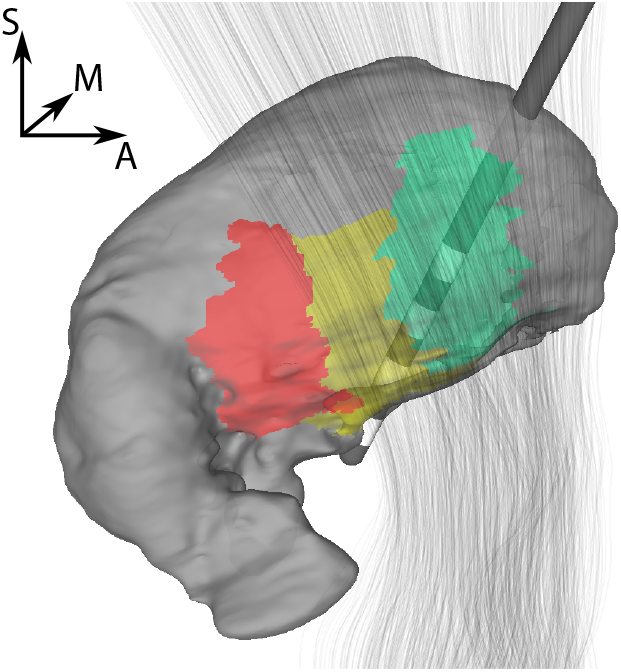
Surface visualization of lead placement relative to thalamus (gray) and IC (gray lines) for P1R. Selected subregions of the thalamus are highlighted (Vc/VPL - red, VIM/VLpv - yellow, Vop/VLa - green).

### 2.4. Atlas registration and fiber tractography

Preoperative imaging of each patient was used to obtain individual segmentations of the target and avoidance regions VIM, Vc, and IC:

The T1w-MRIs were nonlinearly registered to the “MNI ICBM 152 nonlinear 6th Generation Average Brain” (MNI-ICBM152, Grabner et al. 27) using ANTs [6]. For this average brain, surface segmentations of the posteroventral part of the ventrolateral nucleus (VLpv) and the ventral posterolateral nucleus (VPL), which correspond to VIM and Vc, respectively, in Jones nomenclature (see, e.g., Bakay 7, Figure 11.2), were obtained from the Morel atlas [38], which is aligned with the MNI-ICBM152 average brain. Individual surface segmentations of VLpv and VPL aligned with the patient’s T1w-MRI were obtained by transforming the surface segmentations from the MNI-ICBM152 average brain to the patient’s MRI using the nonlinear transform computed in the registration.

Individual tractography of the IC was performed using DSIStudio (dsi-studio.labsolver.org) for each patient and hemisphere [35]. A seed region to generate IC tractography, corresponding to the posterior limb of the IC, was manually segmented based on the fractional anisotropy and fiber orientation (see large solid arrow in Jellison et al. 34, Figure 11.2). We restricted our segmentation to the posterior limb of the IC, since this seed region resulted in fiber tracts that directly passed the stimulation site. Additionally segmenting the anterior limb of the internal capsule resulted in additional fibers more distant to the lead contacts, which therefore did not affect the optimization result, but led to a higher computational effort. To ensure that only superior-inferior fibers, which represent the IC, were included, we added a transversal plane inferior to the thalamus as a region of interest (ROI).

### 2.5. Finite element simulation of DBS

We used the finite element method (FEM) to solve the bioelectric field problem of DBS for the Medtronic 3387 lead [15]. We obtained surface meshes of skin, skull, cerebrospinal fluid (CSF), gray matter, and white matter using SimNIBS 2.1 (http://www.simnibs.de). We localized the lead positions based on the postoperative CT images and generated a triangular surface mesh of the Medtronic 3387 lead with a 0.5mm thick encapsulation layer [30]. Based on these surface meshes, a tetrahedral volume mesh was generated using TetGen [55]. This mesh incorporated the nodes at which we evaluated the electric potential *u* as mesh vertices. These nodes are distributed on a regular grid with a size of 20 × 20 × 30 mm and an internode distance of 0.2mm around the center of the lead. This predefined grid defines the volume Ω on which we perform the optimization, but we removed the nodes inside the lead and the encapsulating tissue. The resulting tetrahedral meshes consisted of about 2.3 million nodes and 13.7 million elements.

We chose the conductivities for the different tissue compartments according to Table 3. For the white matter compartment, we calculated DTI tensors based on the DSI recordings using DTIFIT (https://fsl.fmrib.ox.ac.uk/fsl/fslwiki) and scaled each tensor following the “direct approach with volume constraint” to obtain anisotropic conductivity tensors [59, 29]. We solved the bioelectric field problem with a linear FEM using SCIRun 5 (http://www.sci.utah.edu/cibc-software/scirun.html). We clipped the elements inside the lead and modeled active contacts by imposing a Dirichlet boundary condition set to the stimulation voltage at the contact surface and a homogeneous Neumann boundary condition at the insulating lead shaft. We modeled the return electrode by imposing a Dirichlet boundary condition of 0V where the mesh was cut off in the patient’s neck, as we did not consider bipolar stimulation in this study. The passive contacts were modeled as linked nodes, i.e., considered floating.

**Table 3:** Tissue conductivities

| Tissue | Value [S/m] | Reference |
| --- | --- | --- |
| Skin | 0.43 | Haueisen et al. [31], Ramon et al. [49] |
| Skull | 0.01 | Dannhauer et al. [22] |
| CSF | 1.79 | Baumann et al. [8] |
| GM | 0.33 | Haueisen et al. [31], Ramon et al. [49] |
| WM | 0.14 | Haueisen et al. [31], Ramon et al. [49] |
| Encapsulation | 0.10 | Grill and Mortimer [28] |

Due to the linearity of the problem, it is sufficient to solve the bioelectric field problem for a unit voltage of –1V at each contact while all other contacts are floating. For each solution of the electric potential, we computed the Hessian matrix *H*, which is needed for the optimization algorithm described in Section 2.6, at each node of Ω. The voltage distributions *u*_**c**_ and Hessian matrices *H*_**c**_ for general stimulation voltages **c** = (*c*_0_, *c*_1_, *c*_2_, *c*_3_) follow by linear combination, where the *c_i_* indicate the voltage applied at contact *i*.

### 2.6. Optimization algorithm

To obtain multipolar stimulation configurations, we applied a modification of the optimization algorithm presented by Anderson et al. [4]. The algorithm relies on the activating function as a predictor for neuronal activation [42, 50, 65, 40]. To easily obtain the activating function, we numerically calculated the Hessian matrix of partial second derivatives of the electric potential *u*_**c**_:

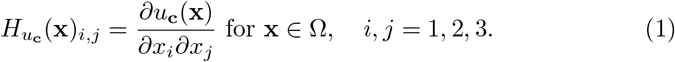

The algorithm presented by Anderson et al. [4] achieves target stimulation by maximizing the sum of the mean of the three eigenvalues of the Hessian taken over all positions in the target region; in avoidance regions the maximal eigenvalue of the Hessian is kept below a threshold *α* (see also Section 2.7). For the targeting or avoidance of fiber tracts, the second spatial derivative in the direction of the fiber tract, which can be computed as *v^t^Hv* for a direction *v*, is considered instead of the eigenvalues. To limit the power output, the maximal charge density at each contact is limited.

Varying the approach of Anderson et al. [4], we used the value of the second derivative of the electric field in a direction perpendicular to the shaft of the DBS electrode to estimate activation of both the target and avoidance regions [14]. This choice corresponds to the assumption of an axon orientation perpendicular to the electrode shaft, as also usually chosen for the multicompartment axon models that are used to compute the volume of tissue activated (VTA) [14, 12, 5]. Based on the Hessian matrix, the activation at a position x with a corresponding direction **n**_*p*_ perpendicular to the electrode shaft can be computed as

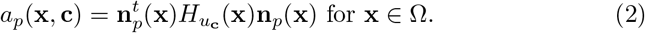

Accordingly, the objective function to measure activation of the target region, Ω_VIM_ ⊂ Ω, is defined as

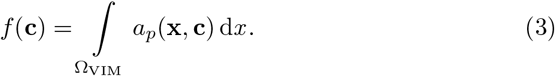

We introduced an additional penalty term to the optimization functional to favor stimulation settings with a higher efficiency by penalizing the stimulation of brain tissue outside the target region Ω_VIM_, also referred to as stimulation spill [19]. This term quadratically penalizes stimulation of the volume Ω – Ω_VIM_:

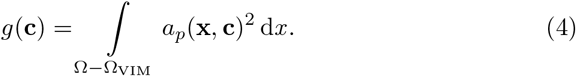

The linear weighting of *a_p_* in *f*, which we aim to maximize, ensures that the mean of the stimulation over the whole target region is maximized and that the value of *f* is not dominated by a few, large outliers of *a_p_*, which could result in a strong stimulation of only small parts of the target volume. The quadratic weighting of *a_p_* in *g*, which we aim to minimize, avoids strong outliers in the stimulation of the nontarget volume.

Stimulation of Vc and IC is avoided by adding constraints to the optimization problem. For the avoidance of the Vc, we kept the value of *a_p_* below the threshold *α* in the region Ω_Vc_. For the avoidance of the IC, we make use of the fiber orientations obtained from tractography. We denote the fiber orientation at position **x** by **n**_*f*_(**x**) and define *a_f_* according to (2) with **n**_*p*_ replaced by **n**_*f*_. As for the avoidance of the Vc, *a_f_* is kept below *α* to avoid stimulation of the IC.

With these constraints, our optimization problem can be formulated as the constrained optimization problem:

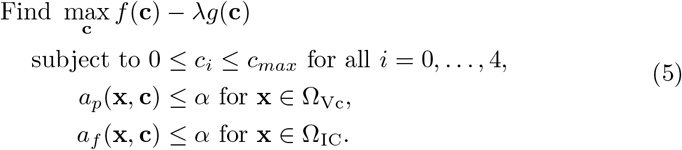

The optimization problem admits a unique solution (a proof can be derived following Wagner et al. 64), resulting in a (multipolar) stimulation pattern **c** = (*c*_0_, *c*_1_, *c*_2_, *c*_3_), with *c_i_* being the stimulation voltage at contact *i*.

The algorithm has been evaluated only for the optimization of (multipolar) cathodic stimulation thus far [4], i.e., all active contacts are concurrently stimulating at different voltages as cathodes, and the IPG serves as the return electrode. In practice, the IPG has to provide independent current/voltage sources for each contact to implement such stimulation patterns. To achieve cathodic stimulation, we enforce 0 ≤ *c_i_* in (5), as we set *c_i_* = –1V in our FEM simulation. *c_max_* allows us to define a general maximal stimulation voltage for safety reasons or, if necessary, also for a single contact, e.g., due to known side effects. This parameter was not employed in our study. The parameter *λ* determines how much weight is given to the efficiency of the stimulation in the optimization. We used *λ* = 0.003 in this study.

In the implementation of the algorithm, the integrals in the expressions for *f* and *g* turn into summations over all (grid) points of Ω. We solved this constrained optimization problem using the Python package cvxopt (cvxopt.org).

### 2.7. Empirical determination of the sensitivity threshold α

In theory, the threshold *α* represents the value of the activating function above which neuron firing occurs as a result of the stimulation, i.e., the firing threshold [4]. Approximate values for this threshold have been derived in simulations of multicompartment neuron models [41, 14, 16, 4]. However, the computation of these thresholds depends on parameters that are not accessible in practice and are often computed under simplified model assumptions, i.e., the axons are assumed to be perfectly straight and perpendicular to the electrode shaft. Furthermore, it is unclear how to link these thresholds to the occurrence of side effects in a computational model. For example, for a representation of the IC through fiber tracts as prepared in Section 2.4, do side effects occur when the firing threshold is exceeded for a single tract, for a certain volume fraction of the tracts, or for all tracts?

Therefore, we do not interpret *α* as a fixed firing threshold, but treat it as a patient-specific sensitivity threshold that has to be derived empirically. Algorithm 1 is an example algorithm to determine *α*.

#### Algorithm 1 Empirical determination of the sensitivity threshold *α*

~~~
*α* = 0
**repeat**
   • *α* = *α* + Δ*α*
   • Solve optimization (5) to obtain stimulation configuration **c**
   • Program patient IPG with stimulation configuration **c**
   • Observe therapeutic and side effects
   **if** Improvement of therapeutic effects **and** no side effects **then**
      **c**_*th*_ = **c**
   **end if**
**until** Side effects observed
Program patient IPG with stimulation configuration **c**_*th*_
~~~

By applying Algorithm 1, the clinical DBS programming decreases to individually adjusting the single parameter *α*, instead of determining therapeutic and side-effect thresholds for each contact. The step width Δ*α* can be individually determined by the clinician based on the increase of the stimulation voltages **c** with increasing *α*, and a first review of a subset of possible configurations can be performed ahead of the programming session. The choice of stimulation frequency and pulse width remains up to the physician. As the firing threshold of axons changes with frequency and pulse width [14, 2], a change of these parameters during a programming session might require adjustements of *α*.

In our retrospective analysis, it was not possible to observe therapeutic and side effects. Instead, we made use of the thresholds that were obtained during the monopolar review (see Section 2.2, Table 2). To obtain optimized stimulation configurations, we incrementally increased the sensitivity threshold α in steps of 0.1 mV/mm^2^ following Algorithm 1 until either the therapeutic or the side-effect threshold was exceeded at one contact. If only a therapeutic threshold was exceeded, the resulting configuration was adopted. If a side-effect threshold was exceeded, α was reduced to the previous value, and the configuration for which all contact voltages fell below the side-effect thresholds was adopted (Table 4).

**Table 4:**
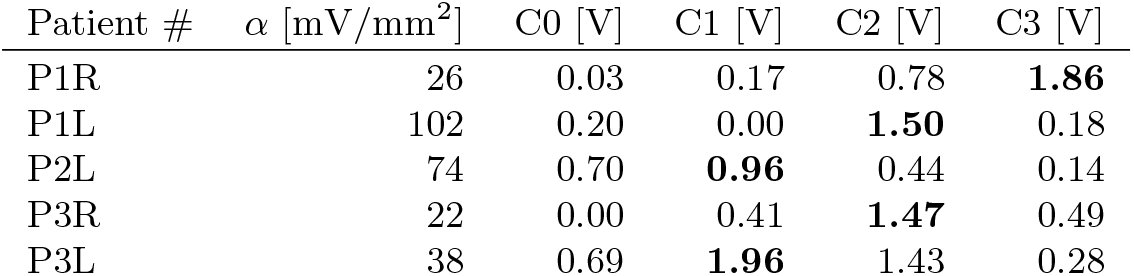
Optimization results (boldfaced type marks clinically selected contact for comparison).

## 3. Results

The optimized stimulation settings obtained following the description in Section 2.7 are shown in Table 4. Most importantly, the optimization algorithm assigns the highest voltage to the contact that was found to drive the best therapeutic effect during the monopolar review in all cases. In the three cases where an alternative therapeutic contact was determined (P1L, P3R, P3L), the optimization algorithm assigned the second highest voltage to these contacts, and for P2L the second highest voltage was assigned to the second contact for which a therapeutic threshold was found. For P1R, a therapeutic threshold was found for only one contact.

To quantitatively compare the optimization results and those of the monopolar review, we computed the predicted target activation and the predicted power consumption. The predicted target activation is a common measure to evaluate the performance of optimization algorithms, and is calculated as the volume fraction of the target region for which the stimulation exceeds a predefined threshold [4, 47, 20]. The predicted target coverage is a computational tool to simulate and compare the efficiency of different stimulation patterns; it does not necessarily correspond to the target activation in practice. We visualize the percentage of the target volume Ω_VIM_ for which *a_p_* is larger than the sensitivity threshold *α* in Figure 2a, where for each patient and hemisphere the individually determined values for *α* are used (Table 4).

**Figure 2:**
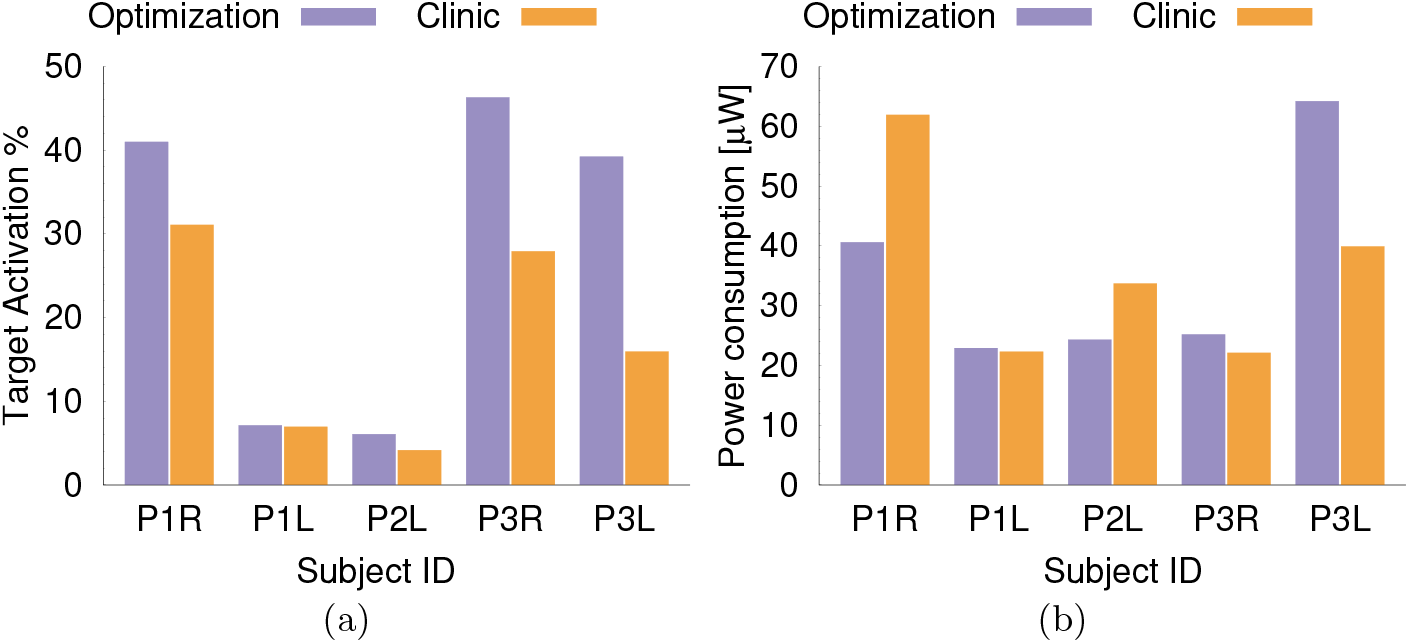
a) Target activation, measured in % of Ω_VIM_ for which *a_p_* is above the sensitivity *α*, for the result of the optimization algorithm (purple) and clinically chosen stimulation setting (orange). b) Predicted power consumption for the result of the optimization algorithm and clinically chosen setting.

We computed the predicted power consumption following Fakhar et al. [26]

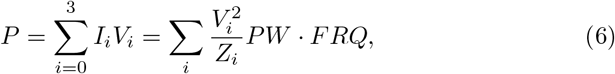

where *I_i_*, *V_i_*, and *Z_i_* are current, voltage, and impedance at contact i, respectively, and PW and FRQ are pulsewidth and frequency. Since we did not have measured impedance values available for each contact, we instead relied on impedance values obtained from the FEM simulations to calculate the predicted power consumption, which is sufficient for our goal of comparing the predicted power consumption of the optimized and the clinically found stimulation settings. The obtained impedance values ranged from 1095 Ω to 1131 Ω, which is within the commonly observed range [15].

Figure 2a and Table 5 show that the optimization result leads to a predicted target activation that is at least equal to the activation of the clinically determined configurations in all cases. For all cases except P1L, the predicted target activation could be clearly improved by at least about 32% (P1R) using the algorithm-derived configurations that employ two or more contacts concurrently. Comparing the changes in predicted target activation with the changes in predicted power consumption (Figure 2b, Table 5), we find that in all of these four cases the improvements in target activation outmatch the change in predicted power consumption by at least 50 percentage points. For P1R and P2L, we find improvements in predicted target stimulation of more than 32% and a simultaneous reduction of the predicted power consumption of more than 28%. This result is achieved by concurrently stimulating at multiple contacts with voltages lower than for the clinically determined monopolar setting. For P3R and P3L, we find improvements in predicted target stimulation of more than 66%, but a simultaneous increase of the predicted power consumption of more than 28%. This result is achieved by concurrently stimulating at multiple contacts, where the voltage at the contact that was found to provide the best therapeutic effect during the monopolar review in the optimized multipolar stimulation pattern is nearly equal to the voltage that was clinically determined.

**Table 5:** Change in predicted target activation and predicted power consumption for optimized stimulation settings in comparison to monopolar stimulation with the best therapeutic effect.

| Patient # | Target activation | Power consumption |
| --- | --- | --- |
| P1R | 31.9% | -34.4% |
| P1L | 2.6% | 2.7% |
| P2L | 45.9% | -27.9% |
| P3R | 65.9% | 14.0% |
| P3L | 146.2% | 60.9% |

Next, we analyze two cases in more detail. We chose P1R exemplaric for the four cases in which multiple contacts are activated concurrently, and we chose P1L as the only case in which only a single contact gets assigned a significant voltage.

### 3.1. Analysis of P1R

For P1R, Figure 3 shows that both C2 and C3 are inside the VIM (green), thus similarly contributing to the target stimulation. However, C2 is closer to the Vc than C3, whereas the IC is relatively distant from both of these contacts. C0 and C1 are both (mostly) outside the VIM, thereby stimulating the VIM less efficiently than C2 and C3. Accordingly, C2 and C3 are are the only active contacts in the multipolar configuration obtained from the optimization, with a higher voltage assigned to C3, which stimulates the Vc less.

**Figure 3:**
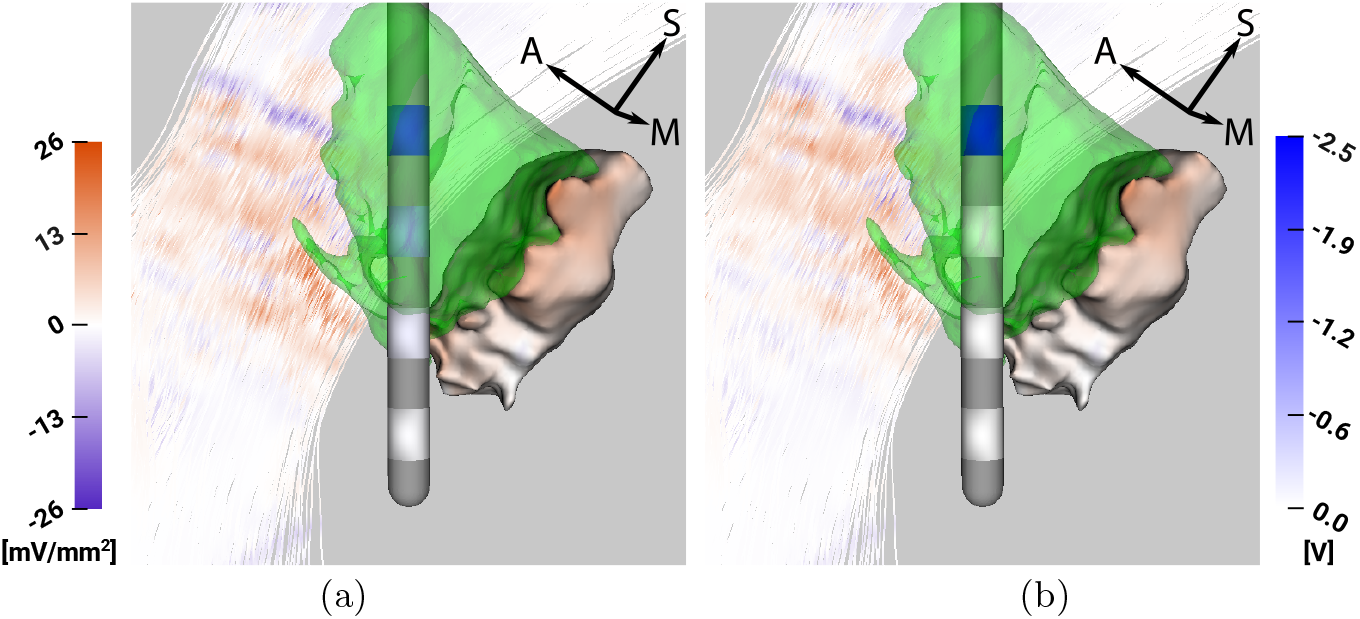
P1R - Visualization of VIM (green), *a_p_* and *a_f_* mapped on Vc and IC (left colorbar in mV/mm^2^) and stimulation voltage mapped on lead contacts (right colorbar in V), respectively, for a) optimization result and b) monopolar stimulation with 2.5 V at C3.

**Figure 4:**
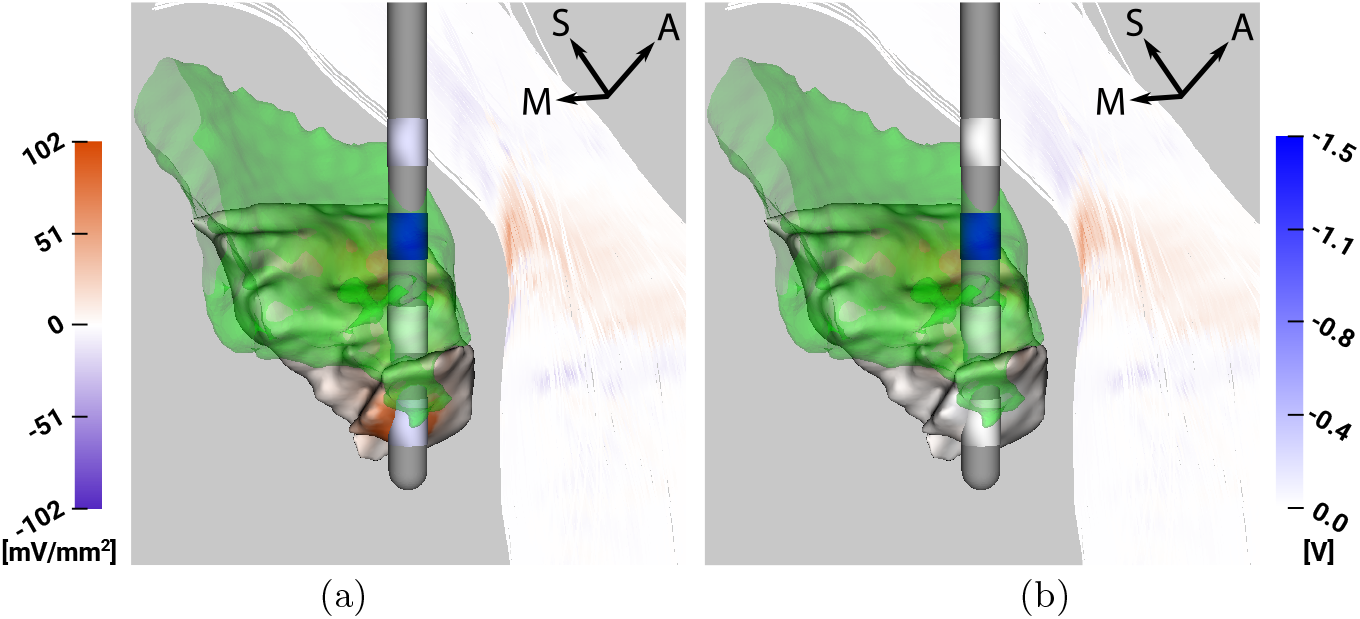
P1L - Visualization of VIM (green), *a_p_* and *a_f_* mapped on Vc and IC (left colorbar in mV/mm^2^) and stimulation voltage mapped on lead contacts (right colorbar in V), respectively, for a) optimization result, b) monopolar stimulation with 1.5 V at C2.

### 3.2. Analysis of P1L

For P1L, we find that C2 is fully contained in the lateral part of the VIM, whereas C1 is found to be inside both the target and avoidance region. C0 lies inferior to the VIM partially inside the avoidance region, and C3 lies superior to the VIM. Thus, C1 and C2 would lead to the best target stimulation, but C1 also leads to a strong stimulation of the Vc. All contacts lead to only a low stimulation of the IC. These considerations explain why the optimization algorithm leads to a nearly exclusive activation of C2 – the other contacts are either in the avoidance region (C0, C1) and/or do not stimulate the VIM efficiently (C0, C3). Already the slight activation of C0 in the optimized stimulation setting leads to noticeable stimulation of the Vc.

## 4. Discussion

In this study, we performed a retrospective evaluation of an algorithm for the automated optimization of DBS configurations that utilize independent current/voltage sources for each contact, as they are available in novel IPGs [1, 58]. The computed stimulation configurations are in good agreement with the results of the monopolar review; in all cases, the optimization algorithm assigned the strongest stimulation voltage to the contact that was clinically selected, and the predicted target activation achieved by the optimization algorithm was at least equal to the clinically determined settings; in four of five cases, the predicted target activation was improved.

### 4.1. Impact

The results obtained in this study are an important demonstration of the capabilities of the applied optimization algorithm, and they encourage its further evaluation for an application in clinical practice. Our results indicate that Vc and IC are appropriate avoidance regions for the automated optimization of VIM stimulation.

To date, most clinicians explore the space of possible stimulation parameters by first evaluating each contact individually in a monopolar review and then using clinical intuition and heuristics to refine their results. This method, since it relies on intuition, is challenging to teach and not guaranteed to find an optimal result. As novel electrodes with an increased number of contacts and IPGs with multiple independent current/voltage sources continue to proliferate, the parameter space for DBS programming grows exponentially, and manual exploration of the parameter space becomes even more complicated and time consuming. Our proposed semiautomated method improves upon this manual process by replacing the review of stimulation voltage or current on each electrode contact by the review of a single parameter, the sensitivity threshold *α*. This reduction of the parameter space is achieved by applying an optimization algorithm that utilizes individual representations of target and avoidance regions obtained from the pre- and postoperative imaging data, which are functionally ignored during a monopolar review. The necessity of iteratively determining *α* (vs. directly computing stimulation settings based on a fixed threshold) arises from our inability to perfectly map models of stimulated tissue to perception (i.e., side effects) at the level of each individual patient, although this limitation may ultimately recede as our modeling and imaging methods improve (see also Section 2.7).

The approach we chose in this study to obtain the sensitivity threshold *α* (Algorithm 1) could be directly applied in clinical practice: The clinician starts to program the patient with a multipolar stimulation configuration that was determined using a low value of *α*. Subsequently, *α* gets incrementally increased, and the patient programming is updated with a new optimization result. This step is repeated until side effects occur, where the initial value of *α* and the step width Δ*α* are determined by the clinician based on the stimulation voltages proposed by the optimization. Finally, the configuration that achieved the best therapeutic effects with the lowest stimulation amplitudes and without side effects is selected. Our simulation results predict that the stimulation configurations obtained from this procedure might lead to equivalent, if not better, target activation compared to the results achieved with a classical monopolar review. In future studies, it has to be determined whether this equivalent/better predicted target activation also leads to equivalent/better therapeutic outcomes.

### 4.2. Related work

Different algorithms for the automated optimization of DBS configurations, which mainly differ in the formulation of the objective function (i.e., (5) in our case), have been proposed. The algorithm applied in this study is based on the work of Anderson et al. [4], and the objective function we employed is comparable to the one proposed by Wagner et al. [64] for transcranial direct current stimulation (tDCS). Through the addition of the functional *g* as defined in (4), we were able to modify this algorithm to simultaneously maximize stimulation of the target region, minimize stimulation of other brain regions, and stay below the side-effect threshold in the avoidance regions.

Similar to our approach, Peña et al. [47] defined an objective function based on a modified activating function (MAF), which is a smoothed activating function. They defined an objective function with three objectives: maximize the volume of the target region that is stimulated above a fixed MAF threshold (MAFT), minimize the volume of the avoidance region that is stimulated above the same MAFT, and minimize power consumption. The weighting factors between these objectives were chosen based on a subjective ranking of the importance of the objectives. Compared to our objective function, the stimulation of the avoidance region is not as strictly enforced in this approach. Whereas our approach directly enforces this avoidance through a constraint in the optimization problem (5), a stimulation above threshold in the avoidance region is penalized in the approach of Peña et al. [47], but not strictly excluded. For example, if the threshold is at the same time exceeded for a large volume of the target region and a clearly smaller volume of the avoidance region, a configuration might still be considered optimal. The limitation of power consumption serves a similar purpose as our functional *g*, enforcing the efficiency of the stimulation.

Instead of using the activating function as a measure of stimulation, Cubo et al. [20] proposed an optimization algorithm based on the electric field strength.

For a given contact configuration and a fixed electric field threshold, this approach penalizes understimulation in the target region quadratically, whereas overstimulation is penalized linearly. Stimulation of avoidance regions exceeding the threshold is prevented by constraining the optimization problem, similarly to (5). Cubo et al. [20] concluded that, in order to achieve meaningful optimization results, the brain volumes to be avoided should be exactly specified and the efficiency of the stimulation should be considered in the optimization.

Furthermore, in line with our considerations in Section 2.7, Cubo et al. [20] concluded that “the threshold values […] to decide which brain volumes are stimulated […] appear to be patient specific.”

The study designs to evaluate the different proposed optimization approaches and the parameters to judge the optimization results differ vastly between the discussed studies by Anderson et al. [4], Peña et al. [47], and Cubo et al. [20].

It is therefore not possible to make any conclusions about which of the proposed algorithms would lead to the best results in practice based on the current knowledge. A comparison of different optimization approaches in a simulation or patient study has yet to be performed. Such a study is not only important to compare the strengths and weaknesses of the different optimization approaches, but merging the evidence from multiple algorithms might also help to increase the trust of clinicians in the results of optimization algorithms.

The proposed algorithms focus on optimizing the stimulation voltages for optimal target activation, but they do not directly take into account stimulation pulse width and frequency, which can also have a significant influence on the stimulation effects [52, 10, 2]. For now, the choice of pulse width and frequency remains the responsibility of the clinician.

In our approach, the balance between optimal stimulation of the target and power efficient stimulation can be regulated through the choice of the parameter *λ*. To keep the application of the optimization algorithm simple, it is important that *λ* does not have to be individually adjusted for each patient. In our study, we kept *λ* fixed at a value of 0.003 for all patients and leads. This choice of *λ* led to optimization results in which the highest voltage was assigned to the contact that was found to lead to the best therapeutic effect in the monopolar review for all patients and leads. Furthermore, the optimized stimulation patterns were predicted to be more efficient than the monopolar stimulation in four of five cases, as the increase in predicted target activation exceeded the variation in predicted power consumption by more than 50 percentage points. However, the choice of *λ* depends on many implicit parameters of the modeling pipeline, so that this value of *λ* is for now valid only for the specific modeling pipeline described in Sections 2.5 and 2.6. In future studies, generally applicable values of *λ* should be derived in larger patient groups, and separate patient groups should be used to determine *λ* and to evaluate the optimization results to avoid any bias.

Alternatively, to directly obtain numerous optimization results for varying weighting parameters, Peña et al. [47] performed a particle swarm optimization and computed a Pareto front. It has to be determined in future studies whether users prefer to obtain a single optimization result for a fixed parameter or numerous optimization results to choose from.

With regard to the sensitivity threshold, values in a range from 5 - 40 mV/mm^2^ as thresholds for axon activation have been reported [50, 40, 14, 39]. For P1R, P3R, and P3L, the empirically determined sensitivity thresholds (see Table 4) are within this range, whereas this range is exceeded for P1L and P2L. The variation of *α* between 20 and over 100 mV/mm^2^ underlines the importance of individually determining *α* to achieve a proper stimulation.

### 4.3. Limitations

Our results are a first indication that automatically optimized stimulation settings for multipolar stimulation can lead to meaningful results in comparison to clinical data, while at the same time improving the predicted target coverage. However, with three patients and five evaluated leads, the sample size of this study is small, and further research is needed to validate the results of this study.

Besides the comparison of the results of the optimization algorithm to those of the monopolar review, we evaluated predicted target activation and predicted power consumption to determine the possible improvements in stimulation. These simulation-based measures can give first hints regarding possible improvements in stimulation through the use of optimized stimulation settings, but they cannot replace a direct evaluation of the optimized stimulation settings in patients. An evaluation in patients has to be a main goal of future studies.

The retrospective study design limits the identification of stimulation settings that possibly cause side effects. For P3L, our approach to determine the sensitivity threshold *α* led to an optimization result that assigned high stimulation voltages to multiple contacts, which led to a major increase in predicted target activation (+146.2%, Table 5) and in predicted power consumption (+60.9%). In practice, despite being below the side-effect threshold for monopolar stimulation for each single contact, stimulation with the optimized configuration might nevertheless lead to side effects. Stimulation in the avoidance regions resulting from multipolar stimulation is the sum of the stimulation from each contact; stimulation with multiple contacts at relatively high voltages in the multipolar setting could lead to a higher stimulation of the avoidance regions than in the monopolar settings for which the side-effect thresholds were determined. This larger amount of stimulation of the avoidance regions through multipolar stimulation settings could not be taken into account in our retrospective study design, where we relied on the side-effect thresholds derived from monopolar stimulation as the stopping criterion for Algorithm 1.

Applying Algorithm 1 for P3L in practice, we would expect to find a smaller value for *α*, since therapeutic and side effects would already be observed before the voltages of the single contacts exceed the thresholds that were observed in the monopolar review. As a result, the theoretically computed improvement of the target activation through the optimized stimulation settings could be reduced, but even at a reduced value of *α* an improved target activation can be expected.

Whereas the motivation for an automatic determination of optimal DBS settings is the emerging use of leads with segmented contacts in combination with IPGs providing multiple independent current/voltage sources in clinical practice, the data utilized in this study were obtained for quadripolar leads with cylindrical contacts. The IPGs used for such leads commonly do not offer multiple independent voltage/current sources, which would be necessary to implement the optimized multipolar stimulation configurations in practice. However, the results of the optimization could be used to guide the clinician in the selection of the relevant contacts during a monopolar review, as the optimization algorithm assigns the highest voltage to the contact that was found to lead to the best therapeutic effect in all cases. Also, the use of double monopolar configurations, with multiple contacts stimulating at the same voltage, might be considered given the optimization results for P1R, P2L, P3R, or P3L.

The optimization problem (5) is less complex to solve for a quadripolar lead instead of a lead with eight or more possibly segmented contacts. Nevertheless, our study is a first demonstration of the feasibility of applying the optimization algorithm in practice. Furthermore, the relatively simple programming of the quadripolar lead used for the patients in this study enabled the clinicians to select close-to-optimal stimulation settings in the monopolar review, helping us to obtain a reliable reference for the evaluation of the optimization results. An equally detailed monopolar review is commonly not performed for leads with segmented contacts due to the time burden for patients.

The reliability of the optimization algorithm depends on the accuracy of the underlying model, i.e., the representations of target and avoidance regions as well as the simulations of the bioelectric fields. In our study, representations of VIM and Vc were obtained by nonlinearly registering highly accurate thalamus segmentations of the Morel atlas [38] to the individual patient MRIs (see Section 2.3). Individual representations of the IC were obtained based on the patient’s DSI, which was processed with high accuracy (see Section 2.1). We manually inspected the quality of the underlying nonlinear registration of the Morel atlas to the patient MRIs and the segmentation of the IC for each patient and found no notable deviations. However, given the low MRI contrast within the thalamus, deviations in the segmentation of VIM and Vc cannot be ruled out and could affect the accuracy of the optimization algorithm. Future studies should investigate how possible inaccuracies in the representation of target and avoidance regions affect the results of the optimization algorithm. At the same time, researchers also need to investigate whether the complex head models used in this study can be simplified without significantly affecting the accuracy of the optimization, e.g., by using simplified volume conductor models that include fewer conductive compartments or by relying on representations of the IC obtained from a brain atlas instead of individual segmentations [32].

Besides geometric inaccuracies, uncertainties in the parameters underlying the individual volume conductor models, e.g., the tissue conductivities, can influence the results of the optimization algorithm. Most conductivity uncertainties affect the impedance of all contacts almost equally, e.g., when an encapsulating tissue compartment with constant conductivity is assumed [15, 54, 21]. Our proposed algorithm is robust against these kinds of uncertainties, because simultaneous changes of the impedance for all contacts lead only to a variation of the sensitivity threshold α, which is in any case determined individually (see Section 2.7, Algorithm 1). Recently, Cubo and Medvedev [21] have presented an approach for the online estimation of tissue conductivities in DBS to decrease the conductivity uncertainties in volume conductor modeling, which could be applied in future studies.

Differences in contact impedances that are not accounted for in the computational model would vary the voltages assigned to the affected contacts in the optimized stimulation configuration. To properly model such impedance differences, more detailed volume conductor models than those commonly used nowadays might be necessary. The conductivities of most compartments, such as gray matter, white matter, or CSF, were shown to have a small influence on the contact impedance [13]. The conductivities of these compartments are often assumed to be homogeneous [54, 19, 21], but even modeling anisotropic white matter conductivities, the contact impedances in our simulations varied only between 1095 Ω to 1131 Ω. Given the strong influence of the encapsulating tissue conductivity on the contact impedance [15], it might therefore be necessary to assign different conductivity values to different segments of the encapsulating tissue to properly model impedance differences.

Accurate impedance values are also important when considering current-controlled stimulation, as provided by most state-of-the-art IPGs. The application of the optimization algorithm for current-controlled stimulation is straightforward using Ohm’s law, but requires knowledge of the different contact impedances.

To minimize computation times, the optimization algorithm used in this study, as well as the algorithms proposed by Peña et al. [47] and Cubo et al. [20], approximate stimulation effects. It was shown that such approximations, whether based on the activating function or on the electric field strength, can properly predict activation spread for monopolar cathodic stimulation when compared to computations of multicompartment neuron models [5, 14]. Furthermore, Peña et al. [47] found good agreement between the activation predicted by the MAF and multicompartment neuron models. A central issue in this regard is the selection of the threshold values for the activating function, as these may depend on multiple parameters [14]. In this study, we circumvented this problem by iteratively determining the sensitivity threshold *α* (see Section 2.7).

Besides the effects of model simplifications (approximation of neuron activation through activating function), the effects of numerical inaccuracies also have to be taken into account. The numerical method that was used, i.e., a linear first-order FEM, was shown to achieve high numerical accuracies for bioelectric field simulations. We took great care in the creation of the mesh and chose a resolution that guarantees an accurate solution of the bioelectric field problem. To maximize the accuracy of the simulation at the points at which the voltage was evaluated, we included all these points in our finite element mesh. In future studies, especially when including novel leads with segmented contacts that lead to more complex electric field patterns, the use of current-preserving FEM approaches should be considered to avoid numerical errors [25, 63]. The use of the MAF instead of the activating function as proposed by Peña et al. [47] also might alleviate numerical inaccuracies.

## 5. Outlook

The results of this retrospective study are an important demonstration of the applicability of the optimization algorithm to automatically determine DBS settings exploiting multipolar settings with multiple independent current/voltage sources in clinical practice. Whereas the results of this study demonstrate that the chosen target and avoidance regions for VIM DBS lead to meaningful optimization results, similar studies have to be performed to determine the correct/necessary regions for other DBS targets, such as the subthalamic nucleus or the globus pallidus internus, and for different lead models, especially those including segmented contacts.

In future studies, the optimization algorithm should be applied during patient programming to evaluate whether the optimized settings lead to better treatment outcomes. The basic approach for an application in practice is laid out in Section 2.7. The results of this study are promising with regard to the simplification of the programming of novel DBS leads with more than four contacts. Furthermore, given that recent simulation studies have demonstrated the possible benefit of anodic stimulation [3], the optimization algorithm should also be evaluated without being restricted to cathodic stimulation in both computational and patient studies.

## Acknowledgements

This work was supported by the National Science Foundation (NSF): US IGNITE 10037840 under Dr. Christopher R. Butson, the Graduate Research Fellowship 1256065 under Daria N. Anderson, and by the Austrian Wissenschaftsfonds (FWF), project I 3790-B27. All data have been obtained within the Neuroscience Initiative project Functional Targeting for Deep Brain Stimulator Placement in Movement Disorders. Christopher R. Butson has served as a consultant for NeuroPace, Advanced Bionics, Boston Scientific, IntelectMedical, Abbott (St. Jude Medical), and Functional Neuromodulation. Christopher R. Butson is also a shareholder of Intelect Medical and is an inventor of several patents related to neuromodulation therapy. All other authors have nothing to declare.

## References

[1] Amon, A. and Alesch, F. [2017]. Systems for deep brain stimulation: review of technical features, Journal of Neural Transmission 124(9): 1083–1091.

[2] Anderson, C. J., Anderson, D. N., Pulst, S. M., Butson, C. R. and Dorval, A. D. [2019]. Neural selectivity, efficiency, and dose equivalence in deep brain stimulation through pulse width tuning and segmented electrodes, bioRxiv. URL: https://www.biorxiv.org/content/early/2019/04/18/613133

[3] Anderson, D. N., Duffley, G., Vorwerk, J., Dorval, A. D. and Butson, C. R. [2019]. Anodic stimulation misunderstood: preferential activation of fiber orientations with anodic waveforms in deep brain stimulation, Journal of neural engineering 16(1): 016026.

[4] Anderson, D. N., Osting, B., Vorwerk, J., Dorval, A. D. and Butson, C. R. [2018]. Optimized programming algorithm for cylindrical and directional deep brain stimulation electrodes, Journal of neural engineering 15(2): 026005.

[5] Åström, M., Diczfalusy, E., Martens, H. and Wårdell, K. [2015]. Relationship between neural activation and electric field distribution during deep brain stimulation, IEEE Transactions on Biomedical Engineering 62(2): 664–672.

[6] Avants, B. B., Tustison, N. and Song, G. [2009]. Advanced normalization tools (ants), Insight j 2: 1–35.

[7] Bakay, R. A. [2009]. Georg Thieme Verlag, Stuttgart, chapter 11 Deep Brain Stimulation for Tremor. URL: http://www.thieme-connect.de/products/ebooks/lookinside/10.1055/b-0034-55961

[8] Baumann, S. B., Wozny, D. R., Kelly, S. K. and Meno, F. M. [1997]. The electrical conductivity of human cerebrospinal fluid at body temperature, IEEE Transactions on Biomedical Engineering 44(3): 220–223.

[9] Benabid, A. L., Pollak, P., Hoffmann, D., Gervason, C., Hommel, M., Perret, J., De Rougemont, J. and Gao, D. [1991]. Long-term suppression of tremor by chronic stimulation of the ventral intermediate thalamic nucleus, The Lancet 337(8738): 403–406.

[10] Brocker, D. T., Swan, B. D., So, R. Q., Turner, D. A., Gross, R. E. and Grill, W. M. [2017]. Optimized temporal pattern of brain stimulation designed by computational evolution, Science translational medicine 9(371): eaah3532.

[11] Buhlmann, J., Hofmann, L., Tass, P. A. and Hauptmann, C. [2011]. Modeling of a segmented electrode for desynchronizing deep brain stimulation, Frontiers in neuroengineering 4: 15.

[12] Butson, C. R., Cooper, S. E., Henderson, J. M. and McIntyre, C. C. [2007]. Patient-specific analysis of the volume of tissue activated during deep brain stimulation, Neuroimage 34(2): 661–670.

[13] Butson, C. R., Maks, C. B. and McIntyre, C. C. [2006]. Sources and effects of electrode impedance during deep brain stimulation, Clinical Neurophysiology 117(2): 447–454.

[14] Butson, C. R. and McIntyre, C. C. [2005a]. Role of electrode design on the volume of tissue activated during deep brain stimulation, Journal of neural engineering 3(1): 1.

[15] Butson, C. R. and McIntyre, C. C. [2005b]. Tissue and electrode capacitance reduce neural activation volumes during deep brain stimulation, Clinical neurophysiology 116(10): 2490–2500.

[16] Chaturvedi, A., Butson, C. R., Lempka, S. F., Cooper, S. E. and McIntyre, C. C. [2010]. Patient-specific models of deep brain stimulation: influence of field model complexity on neural activation predictions, Brain Stimulation: Basic, Translational, and Clinical Research in Neuromodulation 3(2): 65–77.

[17] Contarino, M. F., Bour, L. J., Verhagen, R., Lourens, M. A., de Bie, R. M., van den Munckhof, P. and Schuurman, P. [2014]. Directional steering a novel approach to deep brain stimulation, Neurology 83(13): 1163–1169.

[18] Cubo, R., Åström, M. and Medvedev, A. [2015]. Electric field modeling and spatial control in deep brain stimulation, Decision and Control (CDC), 2015 IEEE 54th Annual Conference on, IEEE, pp. 3846–3851.

[19] Cubo, R., Åström, M. and Medvedev, A. [2016]. Optimization of lead design and electrode configuration in deep brain stimulation, International Journal On Advances in Life Sciences 8: 76–86.

[20] Cubo, R., Fahlström, M., Jiltsova, E., Andersson, H. and Medvedev, A. [2018]. Calculating deep brain stimulation amplitudes and power consumption by constrained optimization, Journal of Neural Engineering.

[21] Cubo, R. and Medvedev, A. [2018]. Online tissue conductivity estimation in deep brain stimulation, IEEE Transactions on Control Systems Technology (99): 1–14.

[22] Dannhauer, M., Lanfer, B., Wolters, C. H. and Knösche, T. R. [2011]. Modeling of the human skull in eeg source analysis, Human brain mapping 32(9): 1383–1399.

[23] Deli, G., Balas, I., Nagy, F., Balazs, E., Janszky, J., Komoly, S. and Kovacs, N. [2011]. Comparison of the efficacy of unipolar and bipolar electrode configuration during subthalamic deep brain stimulation, Parkinsonism & related disorders 17(1): 50–54.

[24] Deuschl, G., Schade-Brittinger, C., Krack, P., Volkmann, J., Schäfer, H., Bötzel, K., Daniels, C., Deutschländer, A., Dillmann, U., Eisner, W. et al. [2006]. A randomized trial of deep-brain stimulation for parkinson’s disease, New England Journal of Medicine 355(9): 896–908.

[25] Engwer, C., Vorwerk, J., Ludewig, J. and Wolters, C. H. [2017]. A discontinuous galerkin method to solve the eeg forward problem using the subtraction approach, SIAM Journal on Scientific Computing 39(1): B138–B164.

[26] Fakhar, K., Hastings, E., Butson, C. R., Foote, K. D., Zeilman, P. and Okun, M. S. [2013]. Management of deep brain stimulator battery failure: battery estimators, charge density, and importance of clinical symptoms, PloS one 8(3): e58665.

[27] Grabner, G., Janke, A. L., Budge, M. M., Smith, D., Pruessner, J. and Collins, D. L. [2006]. Symmetric atlasing and model based segmentation: an application to the hippocampus in older adults, International Conference on Medical Image Computing and Computer-Assisted Intervention, Springer, pp. 58–66.

[28] Grill, W. M. and Mortimer, J. T. [1994]. Electrical properties of implant encapsulation tissue, Annals of biomedical engineering 22(1): 23–33.

[29] Güllmar, D., Haueisen, J. and Reichenbach, J. R. [2010]. Influence of anisotropic electrical conductivity in white matter tissue on the eeg/meg forward and inverse solution. a high-resolution whole head simulation study, Neuroimage 51(1): 145–163.

[30] Haberler, C., Alesch, F., Mazal, P. R., Pilz, P., Jellinger, K., Pinter, M. M., Hainfellner, J. A. and Budka, H. [2000]. No tissue damage by chronic deep brain stimulation in parkinson’s disease, Annals of neurology 48(3): 372–376.

[31] Haueisen, J., Ramon, C., Eiselt, M., Brauer, H. and Nowak, H. [1997]. Influence of tissue resistivities on neuromagnetic fields and electric potentials studied with a finite element model of the head, IEEE Transactions on Biomedical Engineering 44(8): 727–735.

[32] Horn, A., Reich, M., Vorwerk, J., Li, N., Wenzel, G., Fang, Q., Schmitz-Hübsch, T., Nickl, R., Kupsch, A., Volkmann, J. et al. [2017]. Connectivity predicts deep brain stimulation outcome in p arkinson disease, Annals of neurology 82(1): 67–78.

[33] Hunka, K., Suchowersky, O., Wood, S., Derwent, L. and Kiss, Z. H. [2005]. Nursing time to program and assess deep brain stimulators in movement disorder patients, Journal of Neuroscience Nursing 37(4): 204.

[34] Jellison, B. J., Field, A. S., Medow, J., Lazar, M., Salamat, M. S. and Alexander, A. L. [2004]. Diffusion tensor imaging of cerebral white matter: a pictorial review of physics, fiber tract anatomy, and tumor imaging patterns, American Journal of Neuroradiology 25(3): 356–369.

[35] Jiang, H., Van Zijl, P. C., Kim, J., Pearlson, G. D. and Mori, S. [2006]. Dtistudio: resource program for diffusion tensor computation and fiber bundle tracking, Computer methods and programs in biomedicine 81(2): 106–116.

[36] Kirsch, A. D., Hassin-Baer, S., Matthies, C., Volkmann, J. and Steigerwald, F. [2018]. Anodic versus cathodic neurostimulation of the subthalamic nucleus: a randomized-controlled study of acute clinical effects, Parkinsonism & related disorders 55: 61–67.

[37] Koller, W. C., Lyons, K. E., Wilkinson, S. B., Troster, A. I. and Pahwa, R. [2001]. Long-term safety and efficacy of unilateral deep brain stimulation of the thalamus in essential tremor, Movement disorders 16(3): 464–468.

[38] Krauth, A., Blanc, R., Poveda, A., Jeanmonod, D., Morel, A. and Székely, G. [2010]. A mean three-dimensional atlas of the human thalamus: generation from multiple histological data, Neuroimage 49(3): 2053–2062.

[39] Martens, H., Toader, E., Decré, M., Anderson, D., Vetter, R., Kipke, D., Baker, K. B., Johnson, M. D. and Vitek, J. L. [2011]. Spatial steering of deep brain stimulation volumes using a novel lead design, Clinical neurophysiology 122(3): 558–566.

[40] McIntyre, C. C., Mori, S., Sherman, D. L., Thakor, N. V. and Vitek, J. L. [2004]. Electric field and stimulating influence generated by deep brain stimulation of the subthalamic nucleus, Clinical neurophysiology 115(3): 589–595.

[41] McIntyre, C. C., Richardson, A. G. and Grill, W. M. [2002]. Modeling the excitability of mammalian nerve fibers: influence of afterpotentials on the recovery cycle, Journal of neurophysiology 87(2): 995–1006.

[42] McNeal, D. R. [1976]. Analysis of a model for excitation of myelinated nerve, IEEE Transactions on Biomedical Engineering (4): 329–337.

[43] Mohammadi, S., Möller, H. E., Kugel, H., Müller, D. K. and Deppe, M. [2010]. Correcting eddy current and motion effects by affine whole-brain registrations: Evaluation of three-dimensional distortions and comparison with slicewise correction, Magnetic Resonance in Medicine 64(4): 1047–1056.

[44] Ondo, W. G. and Bronte-Stewart, H. [2005]. The north american survey of placement and adjustment strategies for deep brain stimulation, Stereotactic and functional neurosurgery 83(4): 142–147.

[45] Ondo, W., Jankovic, J., Schwartz, K., Almaguer, M. and Simpson, R. [1998]. Unilateral thalamic deep brain stimulation for refractory essential tremor and parkinson’s disease tremor, Neurology 51(4): 1063–1069.

[46] Papavassiliou, E., Rau, G., Heath, S., Abosch, A., Barbaro, N. M., Larson, P. S., Lamborn, K. and Starr, P. A. [2004]. Thalamic deep brain stimulation for essential tremor: relation of lead location to outcome, Neurosurgery 54(5): 1120–1130.

[47] Peña, E., Zhang, S., Deyo, S., Xiao, Y. and Johnson, M. D. [2017]. Particle swarm optimization for programming deep brain stimulation arrays, Journal of neural engineering 14(1): 016014.

[48] Pollo, C., Kaelin-Lang, A., Oertel, M. F., Stieglitz, L., Taub, E., Fuhr, P., Lozano, A. M., Raabe, A. and Schüpbach, M. [2014]. Directional deep brain stimulation: an intraoperative double-blind pilot study, Brain 137(7): 2015–2026.

[49] Ramon, C., Schimpf, P., Haueisen, J., Holmes, M. and Ishimaru, A. [2004]. Role of soft bone, csf and gray matter in eeg simulations, Brain topography 16(4): 245–248.

[50] Rattay, F. [1986]. Analysis of models for external stimulation of axons, IEEE transactions on biomedical engineering (10): 974–977.

[51] Reich, M. M., Brumberg, J., Pozzi, N. G., Marotta, G., Roothans, J., Åström, M., Musacchio, T., Lopiano, L., Lanotte, M., Lehrke, R. et al. [2016]. Progressive gait ataxia following deep brain stimulation for essential tremor: adverse effect or lack of efficacy?, Brain 139(11): 2948–2956.

[52] Reich, M. M., Steigerwald, F., Sawalhe, A. D., Reese, R., Gunalan, K., Johannes, S., Nickl, R., Matthies, C., McIntyre, C. C. and Volkmann, J. [2015]. Short pulse width widens the therapeutic window of subthalamic neurostimulation, Annals of clinical and translational neurology 2(4): 427–432.

[53] Ruthotto, L., Kugel, H., Olesch, J., Fischer, B., Modersitzki, J., Burger, M. and Wolters, C. [2012]. Diffeomorphic susceptibility artifact correction of diffusion-weighted magnetic resonance images, Physics in Medicine & Biology 57(18): 5715.

[54] Schmidt, C., Grant, P., Lowery, M. and van Rienen, U. [2013]. Influence of uncertainties in the material properties of brain tissue on the probabilistic volume of tissue activated, IEEE Transactions on Biomedical Engineering 60(5): 1378–1387.

[55] Si, H. [2015]. Tetgen, a delaunay-based quality tetrahedral mesh generator, ACM Transactions on Mathematical Software (TOMS) 41(2): 11.

[56] Soh, D., ten Brinke, T. R., Lozano, A. M. and Fasano, A. [2019]. Therapeutic window of deep brain stimulation using cathodic monopolar, bipolar, semi-bipolar, and anodic stimulation, Neuromodulation: Technology at the Neural Interface.

[57] Steigerwald, F., Müller, L., Johannes, S., Matthies, C. and Volkmann, J. [2016]. Directional deep brain stimulation of the subthalamic nucleus: a pilot study using a novel neurostimulation device, Movement Disorders 31(8): 1240–1243.

[58] Timmermann, L., Jain, R., Chen, L., Brucke, T., Seijo, F., San Martin, E. S., Haegelen, C., Verin, M., Visser-Vandewalle, V., Barbe, M. T. et al. [2016]. 134 vantage trial: Three-year outcomes of a prospective, multicenter trial evaluating deep brain stimulation with a new multiple-source, constant-current rechargeable system in parkinson disease, Neurosurgery 63: 155.

[59] Tuch, D. S., Wedeen, V. J., Dale, A. M., George, J. S. and Belliveau, J. W. [2001]. Conductivity tensor mapping of the human brain using diffusion tensor mri, Proceedings of the National Academy of Sciences 98(20): 11697–11701.

[60] Van Dijk, K. J., Verhagen, R., Chaturvedi, A., McIntyre, C. C., Bour, L. J., Heida, C. and Veltink, P. H. [2015]. A novel lead design enables selective deep brain stimulation of neural populations in the subthalamic region, Journal of neural engineering 12(4): 046003.

[61] Vitek, J., Jain, R. and Starr, P. [2017]. Intrepid trial: A prospective, double blinded, multi-center randomized controlled trial evaluating deep brain stimulation with a new multiple-source, constant-current rechargeable system in parkinsons disease (p5. 016), Neurology 88(16 Supplement): P5–016.

[62] Volkmann, J., Moro, E. and Pahwa, R. [2006]. Basic algorithms for the programming of deep brain stimulation in parkinson’s disease, Movement disorders: official journal of the Movement Disorder Society 21(S14): S284–S289.

[63] Vorwerk, J., Engwer, C., Pursiainen, S. and Wolters, C. H. [2017]. A mixed finite element method to solve the eeg forward problem, IEEE transactions on medical imaging 36(4): 930–941.

[64] Wagner, S., Burger, M. and Wolters, C. H. [2016]. An optimization approach for well-targeted transcranial direct current stimulation, SIAM Journal on Applied Mathematics 76(6): 2154–2174.

[65] Warman, E. N., Grill, W. M. and Durand, D. [1992]. Modeling the effects of electric fields on nerve fibers: determination of excitation thresholds, IEEE Transactions on Biomedical Engineering 39(12): 1244–1254.

[66] Weaver, F. M., Follett, K., Stern, M., Hur, K., Harris, C., Marks, W. J., Rothlind, J., Sagher, O., Reda, D., Moy, C. S. et al. [2009]. Bilateral deep brain stimulation vs best medical therapy for patients with advanced parkinson disease: a randomized controlled trial, Jama 301(1): 63–73.

[67] Willsie, A. and Dorval, A. [2015]. Fabrication and initial testing of the *μ*dbs: a novel deep brain stimulation electrode with thousands of individually controllable contacts, Biomedical microdevices 17(3): 56.

[68] Xiao, Y., Peña, E. and Johnson, M. D. [2016]. Theoretical optimization of stimulation strategies for a directionally segmented deep brain stimulation electrode array, IEEE Transactions on Biomedical Engineering 63(2): 359–371.

